# Antioxidant defenses of *Francisella tularensis* perturb Aim2 Inflammasome Activation

**DOI:** 10.64898/2026.04.09.717616

**Authors:** Zhuo Ma, Jacob Miller, Kayla Fantone, Chandra Shekhar Bakshi, Meenakshi Malik

## Abstract

*Francisella tularensis* is a Gram-negative bacterium that causes tularemia, a fatal zoonotic disease. *F. tularensis* has been used in the bioweapon programs of several countries. Its potential use as a bioterrorism agent led the CDC to classify *F. tularensis* as a Tier 1 Select Agent. The cytosolic sensor absent in melanoma 2 (Aim2) detects double-stranded DNA in the cytosol of infected cells and subsequently assembles a multiprotein complex known as the inflammasome. Inflammasome activation drives the secretion of IL-1β and IL-18, key proinflammatory cytokines required for controlling *F. tularensis* infection. Prior studies have shown that *F. tularensis* actively suppresses Aim2 inflammasome activation; however, the underlying mechanism remains unknown. We hypothesized that *F. tularensis* suppresses Aim2-mediated responses by modulating the intracellular redox environment. We utilized an *F. tularensis* mutant lacking OxyR (Δ*oxyR*), a transcriptional regulator that controls the expression of major antioxidant enzymes. Our results show that macrophages infected with the Δ*oxyR* mutant exhibit significantly higher levels of Aim2-dependent Caspase-1 and IL-1β than those infected with wild-type bacteria. The expression of interferon regulatory factor 1 and the guanylate-binding proteins GBP2 and GBP5, upstream signaling components of the Aim2 inflammasome, is markedly higher in Δ*oxyR*-infected macrophages than in controls. These changes were absent in Δ*oxyR*-infected NADPH oxidase-deficient macrophages, which are unable to generate reactive oxygen species. Collectively, these findings demonstrate that macrophage redox environment plays a key role in activating signaling components required for Aim2 inflammasome activation. This work advances our understanding of how *F. tularensis*-encoded factors subvert host innate immune defenses.

## INTRODUCTION

*Francisella tularensis* is a Gram-negative bacterium that causes tularemia, a potentially lethal infection (1). Due to its ease of dissemination, high virulence, and remarkable infectivity, *F. tularensis* has historically been used in the bioweapons programs of several countries. Its potential use as a bioterrorism agent led the Centers for Disease Control and Prevention to classify *F. tularensis* subsp. *tularensis* as a Tier 1 Select Agent. *F. tularensis* can be acquired through direct contact with infected animals, tick bites, or consumption of contaminated food products. The most common and lethal form of the infection occurs through direct inhalation of as few as 10 colony-forming units (CFU), which can cause severe pneumonic tularemia (2). *Francisella* infects a wide range of hosts, from arthropods to mammals. *Francisella* includes two species: *F. tularensis* and *F. philomiragia*. *F. tularensis* is further divided into the subspecies *tularensis* (Type A), *holarctica* (Type B), and *mediasiatica*. Type A strains are most common in North America, whereas Type B strains occur in North America, Europe, and Asia. *F. tularensis* subsp. *tularensis* is further subdivided into two clades, AI and AII, with the AI clade predominating in the eastern United States and the AII clade in the western United States. Subspecies *holarctica* is divided into clades BI-BV (3). *F. novicida* and *F. philomiragia* rarely cause disease in healthy individuals; *F. novicida*, though genetically close to *F. tularensis*, remains a distinct species and is widely used as a research model (4). *F. philomiragia*, associated with severe infections in near-drowning cases, occupies an aquatic niche distinct from other *Francisella* species.

During macrophage oxidative burst, host-derived reactive oxygen species (ROS) target invading bacteria. *F. tularensis* counters oxidative stress by inducing multiple antioxidant enzymes essential for its survival. These include superoxide dismutases (SodB and SodC), catalase (KatG), and alkyl hydroperoxide reductase (AhpC) (5–7). SodB (FeSOD) and SodC (CuZnSOD) convert toxic superoxide radicals (O₂⁻) into oxygen and hydrogen peroxide (H₂O₂). KatG then detoxifies H₂O₂ into water or, in the presence of ferrous iron, into hydroxyl radicals. H₂O₂ accumulation also activates the transcriptional regulator OxyR, which induces AhpC, KatG, and Fur, thereby reducing intracellular ferrous iron and limiting hydroxyl radical formation (8). KatG additionally helps neutralize reactive nitrogen species. Meanwhile, AhpC degrades peroxynitrite (ONOO⁻), formed from NO and superoxide, into the less harmful nitrogen dioxide (NO₂⁻) (7, 9, 10). Because OxyR regulates numerous oxidative stress-related genes, it is vital for maintaining redox homeostasis and, thereby, intracellular survival.

During *F. novicida* infection, a small amount of bacterial DNA is released into the cytosol of murine macrophages via bacterial lysis, where it is detected by cGAS, IFI204, and STING, thereby inducing type I interferon (IFN) production (11). Type I IFN signals through its receptor to activate IRF3, which, together with STAT1, STAT2, and IRF9, drives expression of IFN-stimulated genes (ISGs) (12, 13). IFN signaling also induces IRF1, which upregulates guanylate-binding proteins (GBPs) such as GBP2 and GBP5, which lyse bacterial cells, releasing more DNA to fully activate the Aim2 inflammasome (14). Activated Aim2 promotes cleavage of pro-caspase-1 into active caspase-1, which processes IL-1β, IL-18, and gasdermin D (GSDMD). GSDMD pores allow IL-1β/IL-18 secretion, potassium efflux, and pyroptosis. Potassium efflux feeds back to suppress the cGAS-STING pathway, preventing excessive IFN-β, which would otherwise be detrimental. Macrophages lacking *Gsdmd* or *Aim2* show minimal IL-1β/IL-18, do not undergo pyroptosis, and produce abnormally high IFN-β, demonstrating the coordinated protective function of Gsdmd and Aim2 (15). The activation of Aim2 in *F. novicida* infection also requires reactive oxygen species (ROS) (16).

Unlike *F. novicida*, *F. tularensis* suppresses Aim2 inflammasome activation through an incompletely understood mechanism (17, 18). Given that *F. tularensis* strains possess strong antioxidant defense mechanisms, we hypothesized that bacterial modulation of the intracellular redox environment prevents bacterial DNA release and Aim2 activation. Using an *F. tularensis* Δ*oxyR* mutant and strains lacking the antioxidant enzymes SodB, SodC, and KatG, we demonstrate that *F. tularensis* neutralizes macrophage-derived reactive oxygen species to suppress redox-dependent Aim2 inflammasome activation and pro-inflammatory cytokine production. These findings identify the bacterial antioxidant system as a critical mechanism for evasion of host innate immunity.

## MATERIALS AND METHODS

### Francisella Strains

*Francisella tularensis* subspecies *holarctica*, Live Vaccine Strain (LVS) (American Type Culture Collection, Rockville, MD Cat. #29684), obtained from BEI Resources, was used for all experiments. Mutants of *F. tularensis* LVS having a gene deletion in the transcriptional regulator of oxidative stress (Δ*oxyR*), a double mutant carrying a gene deletion of *sodC* and a point mutation in the *sodB* gene (*sodB*Δ*sodC*), the catalase gene deletion mutant (Δ*katG*), and a transposon insertion mutant of the *emrA1* gene (*emrA1*) that have been previously generated and characterized in our previous studies were also used (5, 8, 19, 20). *F. tularensis* LVS and the mutant strains were grown on Mueller-Hinton (MH) chocolate agar plates (Hardy Diagnostics, Santa Maria, CA) containing GC agar base with 1% bovine hemoglobin, 2% v/v IsoVitaleX (BD Biosciences, San Jose, CA), and coenzyme supplements or in Mueller-Hinton Broth (MHB) (BD Biosciences, San Jose, CA) containing beef extract, acid digest of casein, starch, 0.0138% w/v anhydrous CaCl_2_, 0.021% w/v MgCl_2_ x 6H_2_O, 1% v/v 10% w/v glucose) and was supplemented with 2.5% w/v ferric pyrophosphate and 2% v/v IsoVitaleX. Stock cultures were prepared by growing the *Francisella* strains in MHB to an OD_600_ of 0.2, aliquoting into 1 mL tubes, snap-freezing in liquid nitrogen, and storing at -80 °C until further use.

### Generation of Bone Marrow-Derived Macrophages (BMDMs)

BMDMs were derived from wild-type C57BL/6J male and female mice, aged 6-8 weeks and genetically altered mice, including absent in melanoma 2 deficient (*Aim2*^-/-^), nicotinamide adenine dinucleotide phosphate (NADPH) Oxidase deficient mice (*gp91phox*^-/-^) and Gasdermin D (*Gsdmd*^-/-^) mice (Jackson Laboratories, Bar Harbor, ME). Dr. Penghua Wang at the University of Connecticut donated femurs from *Sting*^-/-^ mice, which were used to generate *Sting^-/-^* BMDMs. The mice were euthanized using CO_2_ and cervical dislocation. The femur, tibia, and fibula were excised using sterile instruments to extract and separate the bone marrow cells. The separated cells were then suspended in 20.0 mL of BMDM media containing DMEM, 20% v/v HI-FBS, 30% v/v L-Cell conditioned media, 0.1% v/v 100x Glutamax, 0.1% v/v 100x sodium pyruvate, and 1% v/v HEPES and added to 150 mm tissue culture petri dishes and subsequently incubated at 37°C and 5% CO_2_. To separate the BMDMs from non-monocytic cells, supernatants were transferred into 150-mm tissue culture petri dishes the following day. The monocytes were allowed to grow, proliferate, and differentiate for 6-9 days to form a monolayer. Two to five mL of fresh L-Cell-conditioned media was added every 2-4 days to support the growth. Once high confluency was achieved, the remaining medium was removed, 10.0 mL of cold phosphate-buffered saline was added, and the mixture was placed at 4°C for 10 minutes. The cells were scraped and centrifuged at 1,200 RPM for 10 minutes. Pelleted cells were resuspended in DMEM without any antibiotics and counted using a hemocytometer and 0.4% Trypan Blue stain. For western blot analysis, BMDMs were seeded at a concentration of 5× 10^6^ cells/mL in 12-well tissue culture plates.

### Infection Protocol

For western blot analysis, BMDMs were seeded in a 12-well tissue culture plate at a density of 5 × 10^6^ cells per well. The tissue culture plates were incubated for 16-24 hours at 37°C and 5% CO_2_, allowing the cells to adhere and acclimate. Subsequently, the BMDMs were infected with either *F. tularensis* LVS or the mutant strains at a multiplicity of infection (MOI) of 50. Uninfected BMDMs served as negative controls. The 12-well plates were centrifuged at 1,200 RPM for 5 minutes to synchronize the infection and incubated at 37°C and 5% CO_2_ for 24 hours. The following day, the cell supernatants were removed from each well, placed in 2.0 mL microcentrifuge tubes, and the cells were scraped from the plates and centrifuged at 17,500 RPM for 10 minutes at 4°C to pellet cells. The cell pellets were resuspended in 200.0 μL of Higgins cell lysis/tissue Homogenization and immunoprecipitation buffer (10mM Tris-HCl, pH 7.5, 0.1% sodium dodecyl sulfate (SDS), 0.5% w/v sodium deoxycholate, 0.5% w/v Triton X-100, 1x protease and phosphatase inhibitor cocktail, 0.5mM EDTA, and 0.1mM Dithiothreitol (DTT). These samples were immediately sonicated for approximately 15 seconds each. Cell lysates were stored at -80°C until further use.

### Chemical Inhibitors

Chemical inhibitors Rotenone and Diphenyleneiodonium chloride (DPI) were used. Rotenone (Abcam), diluted in DMSO to 100 mM, is a potent inhibitor of NADH oxidation by inhibiting mitochondrial electron transport chain complex 1; therefore, it stimulates ROS production by mitochondria. DPI (Abcam), diluted in DMSO to 10 mM, is a potent inhibitor of NO synthase, NADPH oxidase, and NADPH cytochrome P450 oxidoreductase, and inhibits mitochondrial ROS. The BMDMs were seeded at 1x10^6^ cells/mL in 12-well plates for 24 hours as previously described. After 24 hours, the cells were treated with inhibitors for four hours. Rotenone and DPI were diluted to 1µM/mL in 10 mL of BMDM-specific media without antibiotics. The medium from the seeded cells was removed and replaced with 1 mL of medium containing the inhibitor. The cells were then incubated for 4 hours at 37^°^C in a 5% CO_2_ atmosphere. After the 4-hour incubation, the medium containing the chemical inhibitor was removed. The cells were then infected at an MOI of 100 with the *F. tularensis* LVS and the antioxidant mutants. Once the bacteria were added to the cells, the chemical inhibitor solution was reapplied, and the plates were centrifuged at 1000 rpm at 4°C for 5 minutes. The cells were then incubated for 24 hours at 37^°^C in the presence of 5% CO_2_ and prepared for western blot analysis as described above.

### SDS-PAGE and Western Blotting

Mini-PROTEAN TGX Stain-Free polyacrylamide gels or Mini-PROTEAN TGX Stain-Free polyacrylamide gels (Bio-Rad) were used for SDS-PAGE. After the proteins were completely separated, the gels were removed and placed in 1x transfer buffer (0.3% w/v Tris, 1.44% w/v Glycine, 20% v/v Methanol) and rocked at room temperature for 10 minutes. Polyvinylidene fluoride (0.2 μm) membranes (Bio-Rad) were activated in >99.8% methanol, washed in ddH_2_O for 1 minute, placed in 1x transfer buffer for 5 minutes, and then western transfer was performed. Once completed, the membranes were removed and washed with H_2_O for 5 minutes. The membranes were then transferred to tubs containing 5% blocking buffer (5% w/v Nonfat Instant dry milk, 10 mM Tris, 100 mM NaCl, 0.05% w/v Tween-20, pH 7.5) and incubated overnight at 4.0 °C while rocking. All primary and secondary antibodies used in this study are shown in Tables 1 and 2. The blots were developed and imaged. Band intensities were quantified using ImageJ.

**Table 1:** List of primary antibodies.

| Antibody Target | Antibody Used | Vendor | Dilution Used |
| --- | --- | --- | --- |
| β-Actin | Monoclonal Anti-β-Actin-Peroxidase Antibody Produced in Mouse | Sigma-Aldrich | 1:20,000 |
| GAPDH |  |  |  |
| IL-1β | IL-1β Rabbit Antibody (H-153) | Santa Cruz | 1:2,000 |
| Caspase-1 | Anti-Caspase-1 mAb (p20) | AndipoGen | 1:3,000 |
| GBP2 | GBP2 Rabbit Polyclonal Antibody | Abcam | 1:1,000 |
| GBP5 | GBP5 Rabbit Polyclonal Antibody | Proteintech | 1:5,000 |
| IRF1 | IRF1 Rabbit Antibody | Novus | 1:1,000 |
| pSTAT1 | pSTAT1 (A2) Monoclonal Antibody | Santa Cruz | 1:500 |
| STAT1 | STAT1 p91 Goat Polyclonal Antibody | Novus | 1:500 |

**Table 2:** List of secondary antibodies.

| Antibody Description | Vendor | Dilution Used |
| --- | --- | --- |
| Mouse Anti-Goat IgG HRP | Santa Cruz Biotechnology | 1:5,000 |
| Mouse Anti-Rabbit IgG HRP | Santa Cruz Biotechnology | 1:5,000 |
| Mouse [m]-IgGκ BP-HRP | Santa Cruz Biotechnology | 1:5,000 |
| Anti-Biotin HRP-Linked Antibody | Cell Signaling Technology | 1:10,000 |

### Statistical Analysis

All data were analyzed using GraphPad Prism software. The statistical significance among sample groups was determined by one-way or two-way analysis of variance (ANOVA), followed by Tukey-Kramer Multiple Comparisons tests. Statistical significance was denoted by any *p*-values of <0.05, and results were expressed as means ± standard error of the mean (SEM) or standard deviation (SD).

## RESULTS

### Enhanced levels of Aim2-dependent bioactive Caspase-1 and IL-1β are observed in BMDMs infected with the Δ*oxyR* mutant of *F. tularensis*

Activation of the inflammasome is associated with proteolytic cleavage of pro-caspase-1 to bioactive caspase-1. The bioactive caspase-1 then cleaves pro-IL-1β into its mature, secreted form. Thus, the detection of bioactive caspase-1 and IL-1β indicates inflammasome activation. We infected BMDMs derived from the wild-type and *Aim2^-/-^* mice with *F. tularensis* LVS and the Δ*oxyR* mutant. The macrophages were lysed and analyzed by SDS-PAGE and western blot to detect bioactive caspase-1 and IL-1β. Significantly higher levels of bioactive caspase-1 and IL-1β were detected in BMDMs derived from wild-type mice infected with the Δ*oxyR* mutant (Fig. 1A and B). However, no detectable or very faint bands of bioactive caspase-1 or IL-1β were observed in *Aim2^-/-^* BMDMs infected either with *F. tularensis* LVS or the Δ*oxyR* mutant (Fig. 1C and D). These results indicated that loss of OxyR leads to enhanced inflammasome activation in wild-type BMDMs infected with the Δ*oxyR* mutant of *F. tularensis* LVS. Furthermore, the higher levels of bioactive caspase-1 and IL-1β in BMDMs infected with the Δ*oxyR* mutant depend on activation of the Aim2 inflammasome.

**FIG 1:**
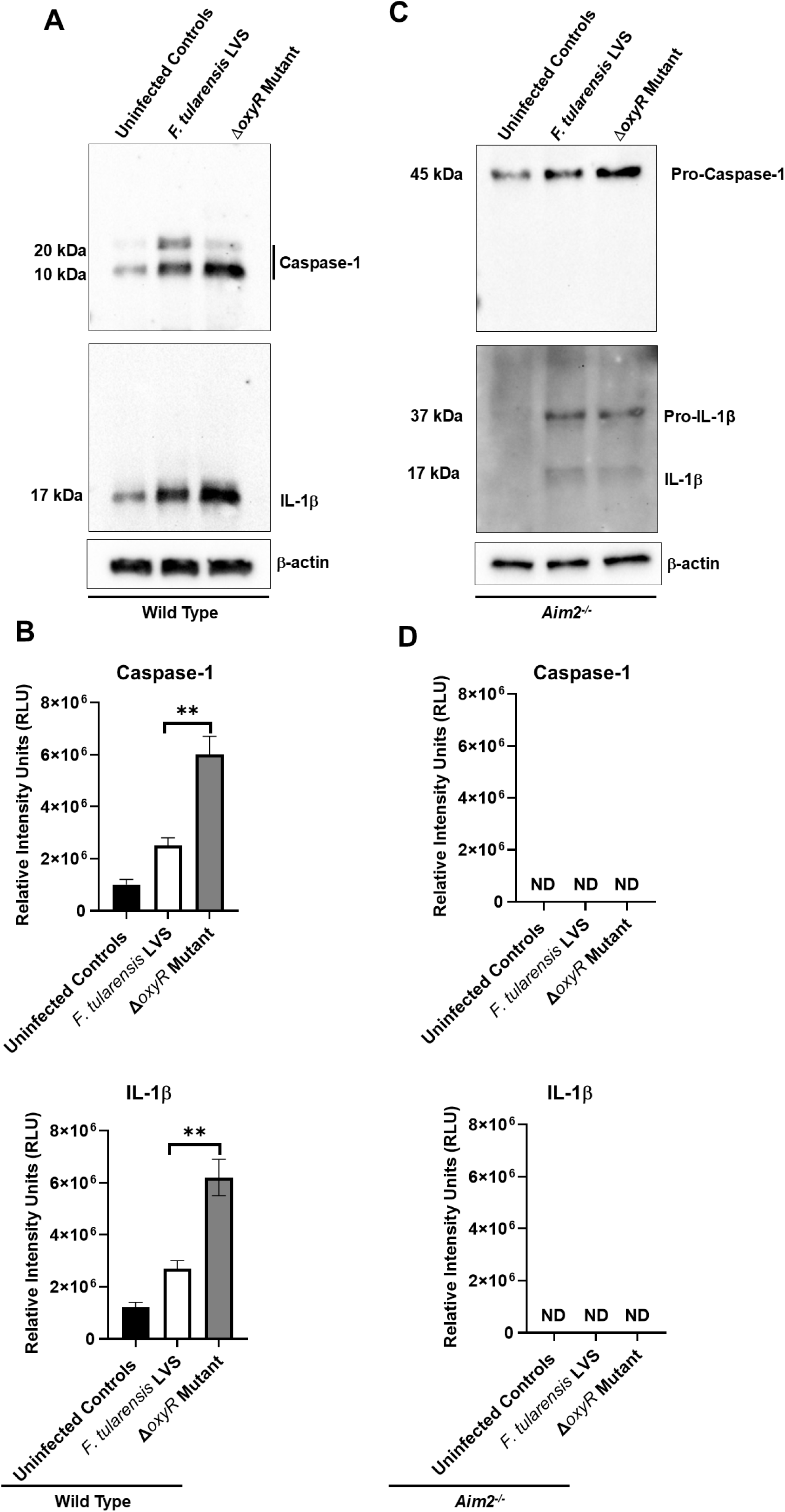
Enhanced levels of bioactive Caspase-1 and IL-1β observed in BMDMs infected with the Δ*oxyR* mutant are Aim2-dependent. BMDMs derived from wild type **(A, B)** or the *Aim2^-/-^*mice (**C, D**) were infected with *Francisella tularensis* LVS or the Δ*oxyR* mutant at a multiplicity of infection of 50 for 24 hours. (**A, C**) Levels of bioactive caspase-1 and IL-1β were assessed by SDS-PAGE followed by Western blot analysis. β-Actin was used as a loading control. Representative immunoblots from three independent experiments are shown. **(B, D)** Quantification of bioactive caspase-1 and IL-1β levels. Band intensities were determined by densitometric analysis, and data from three independent experiments were combined and analyzed using one-way ANOVA. *P* values are indicated (\*\**P* < 0.01).

### Enhanced IRF1 expression and Aim2 inflammasome activation observed in BMDMs infected with the Δ*oxyR* mutant are independent of STING

STING is a DNA sensor that detects small amounts of released DNA and induces type I IFN production (21, 22). Type I IFN binds to its receptor outside the cell and activates IRF3 in an autocrine fashion (23). IRF3 is regulated by transcription factors STAT1, STAT2, and IRF9, which drive the transcription of ISGs (15). Type I IFN then induces the expression of the transcription factor IRF1, which upregulates guanylate-binding proteins (GBPs) such as GBP2 and GBP5, which lyse bacterial cells, releasing more DNA to fully activate the Aim2 inflammasome (14). We next investigated whether the enhanced Aim2 inflammasome activation observed in BMDMs infected with the Δ*oxyR* mutant is due to increased IRF1 expression and whether this expression requires STING. We infected BMDMs derived from the wild-type C57BL/6 and *Sting^-/-^* mice with *F. tularensis* LVS and the Δ*oxyR* mutant. The macrophages were lysed and analyzed by Western blot to determine IRF1 levels. Higher levels of IRF1 were detected in BMDMs derived from wild-type mice and infected with the Δ*oxyR* mutant than those observed for BMDMs infected with *F. tularensis* LVS. Furthermore, similarly elevated levels of IRF1 were observed in *Sting^-/-^* BMDMs infected with the Δ*oxyR* mutant or *F. tularensis* LVS (Fig. 2A and B). These results indicate that enhanced Aim2 inflammasome activation in BMDMs infected with the Δ*oxyR* mutant is due to increased IRF1 expression, which is independent of STING.

**FIG 2:**
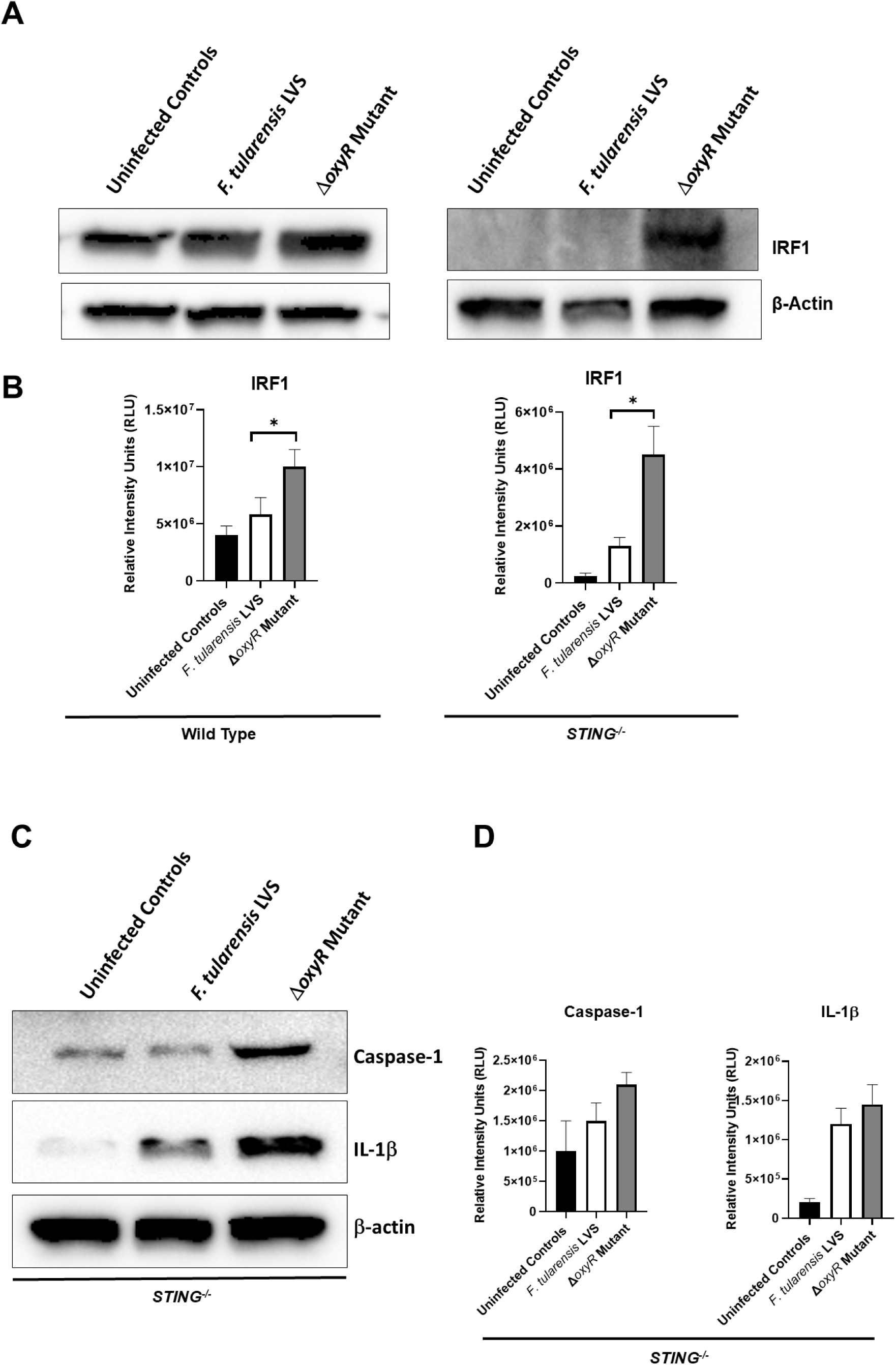
Enhanced interferon regulatory factor 1 (IRF1) expression and Aim2 inflammasome activation observed in BMDMs infected with the Δ*oxyR* mutant are independent of STING. BMDMs derived from wild-type or the *Sting^-/-^* mice were infected with *Francisella tularensis* LVS or the Δ*oxyR* mutant at a multiplicity of infection of 50 for 24 hours. (**A**) Levels of IRF1were assessed by SDS-PAGE followed by Western blot analysis. β-Actin was used as a loading control. Representative immunoblots from three independent experiments are shown. **(B)** Quantification of IRF1 levels. **(C)** BMDMs derived from the *Sting^-/-^* mice infected with *F. tularensis* LVS or the Δ*oxyR* mutant at a multiplicity of infection of 50 for 24 hours. Levels of bioactive caspase-1 and IL-1β were assessed by SDS-PAGE followed by Western blot analysis. β-Actin was used as a loading control. Representative immunoblots from three independent experiments are shown. **(D)** Quantification of bioactive caspase-1 and IL-1β levels. Band intensities (**B and D**) were determined by densitometric analysis, and data from three independent experiments were combined and analyzed using one-way ANOVA. *P* values are indicated (\**P* < 0.05).

We next investigated whether the enhanced IRF1 expression in *Sting^-/-^*BMDMs infected with the Δ*oxyR* mutant is also associated with elevated levels of bioactive caspase-1 and IL-1β. We infected BMDMs derived from the wild-type C57BL/6 and *Sting^-/-^*mice with *F. tularensis* LVS and the Δ*oxyR* mutant. The macrophages were lysed and analyzed by western blotting to detect bioactive caspase-1 and IL-1β. Elevated levels of bioactive caspase-1 and IL-1β were observed in *Sting*^-/-^ BMDMs infected with the Δ*oxyR* mutant than those infected with *F. tularensis* LVS (Fig. 2C and D). Collectively, these results demonstrate that the enhanced levels of IRF1 and Aim2-dependent bioactive caspase-1 and IL-1β in BMDMs infected with the Δ*oxyR* mutant are independent of STING.

### Infection of BMDMs with the Δ*oxyR* mutant results in ROS-dependent activation of STAT1 and elevated IRF1 expression

Our results showed that the enhanced activation of the Aim2 inflammasome in BMDMs infected with the Δ*oxyR* mutant is independent of STING, yet we still observed enhanced IRF1 expression. We hypothesized that IRF1 expression is mediated by STAT1 activation. STAT1 is activated by phosphorylation, followed by its translocation to the nucleus, where it activates transcription of several genes, including IRF1 and GBPs (24). Also, in the nucleus, phosphorylated STAT1 is rapidly degraded by STAT3, making it very difficult to detect activated STAT1 (25). We investigated whether STAT1 mediates the ROS-dependent enhancement of IRF1 expression observed in BMDMs infected with the Δ*oxyR* mutant. We infected BMDMs derived from wild-type and *gp91phox^-/-^* mice with *F. tularensis* LVS and the Δ*oxyR* mutant. The *gp91phox^-/-^* BMDMs cannot produce ROS. The macrophages were lysed and analyzed by SDS-PAGE and Western blot to detect total and phosphorylated STAT1 levels. Identical levels of STAT1 were detected in uninfected and BMDMs infected with the *F. tularensis* LVS or the Δ*oxyR* mutant (Fig. 3A). However, no detectable bands of phosphorylated STAT1 were observed in BMDMs infected with the Δ*oxyR* mutant. Corresponding with the activation of STAT1, significantly enhanced levels of IRF1 were also observed in BMDMs infected with the Δ*oxyR* mutant *F. tularensis* LVS (Fig. 3A and B). Identical faint bands of phosphorylated STAT1 were observed in *gp91phox^-/-^* BMDMs, infected with the Δ*oxyR* mutant *F. tularensis* LVS, indicating a reduced activation of STAT1 in BMDMs infected with the Δ*oxyR* mutant due to the lack of ROS production (Fig. 3C) and correspondingly, the levels of IRF1diminished in Δ*oxyR* mutant-infected *gp91phox^-/-^* BMDMs (Fig. 3C and 3D). These results indicated enhanced ROS-dependent STAT1 activation in BMDMs infected with Δ*oxyR,* leading to elevated IRF1 levels.

**FIG 3:**
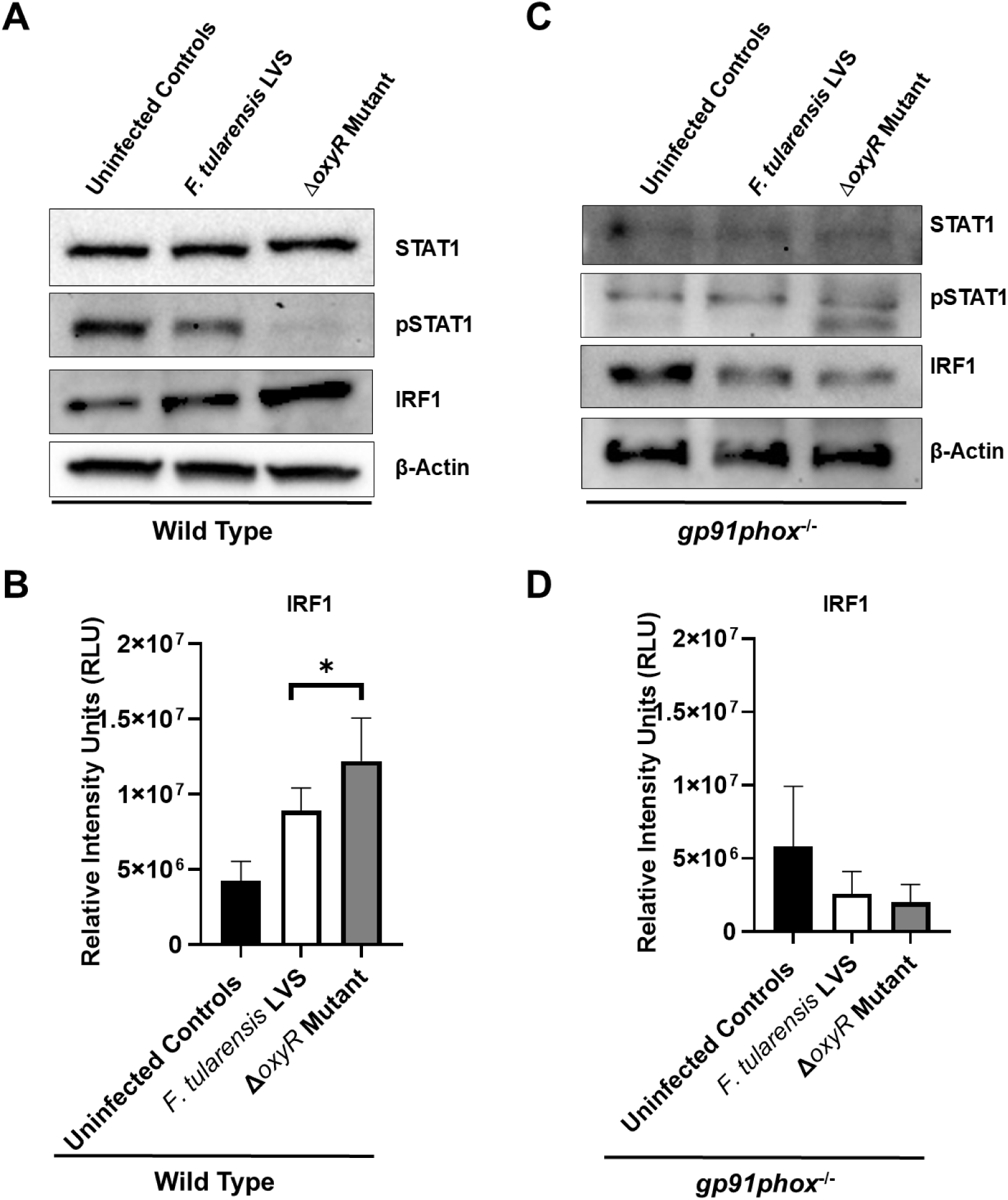
Infection of BMDMs with the Δ*oxyR* mutant results in ROS-dependent activation of STAT1 and elevated IRF1 expression. BMDMs derived from wild-type or the *gp91phox^-/-^* mice were infected with *Francisella tularensis* LVS or the Δ*oxyR* mutant at a multiplicity of infection of 50 for 24 hours. (**A, B**) Levels of STAT 1, pSTAT1, and IRF1in wild type and (**C, D**) in *gp91phox^-/-^* BMDMs were assessed by SDS-PAGE followed by Western blot analysis. β-Actin was used as a loading control. Representative immunoblots from three independent experiments are shown. Band intensities (**B and D**) were determined by densitometric analysis, and data from three independent experiments were combined and analyzed using one-way ANOVA. *P* values are indicated (\**P* < 0.05).

### Enhanced levels of GBP2 and GBP5 observed in BMDMs infected with the Δ*oxyR* mutant are ROS-dependent

Having observed ROS-dependent expression of IRF1, we further investigated whether there is enhanced expression of GBP2 and GBP5 in BMDMs infected with the Δ*oxyR* mutant, and whether this is ROS-dependent. We infected BMDMs derived from wild-type and *gp91phox^-/-^*mice with *F. tularensis* LVS and the *ΔoxyR* mutant. The macrophages were lysed and analyzed by SDS-PAGE and western blot to detect GBP2 and GBP5. Enhanced levels of both GBP2 and GBP5 were detected in wild-type BMDMs infected with the Δ*oxyR* mutant compared to *F. tularensis* LVS-infected BMDMs (Fig. 4A and B). However, these levels diminished and became similar in both *F. tularensis* LVS- and Δ*oxyR*-mutant-infected *gp91phox*^-/-^ BMDMs, indicating reduced expression of GBP2 and GBP5 in BMDMs deficient in ROS production (Fig. 4C and D). These results indicated enhanced ROS-dependent expression of GBPs in BMDMs infected with the Δ*oxyR* mutant.

**FIG 4:**
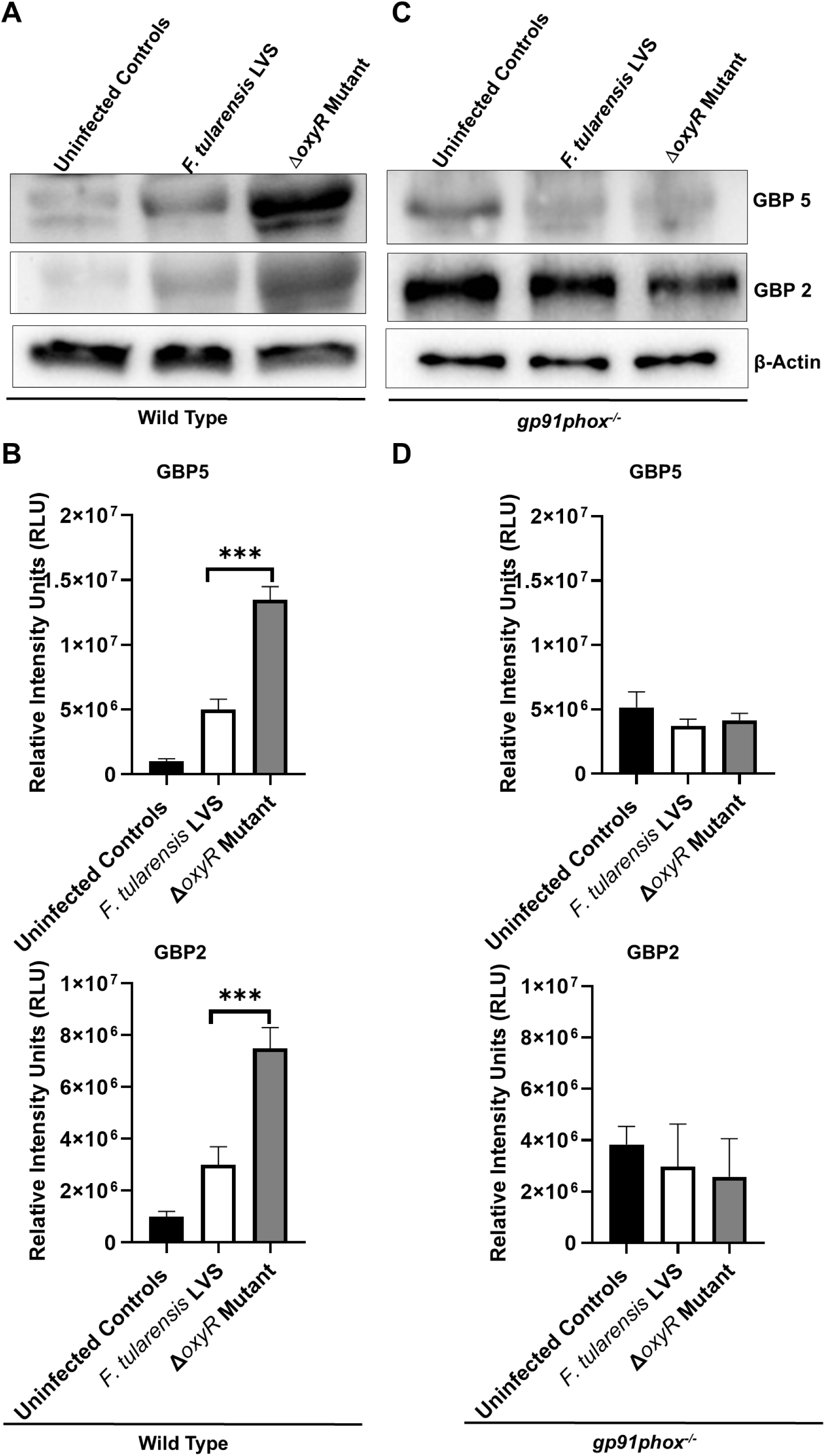
Enhanced levels of guanylate-binding proteins 2 and 5 (GBP2 and GBP5) observed in BMDMs infected with the ΔoxyR mutant are ROS-dependent. BMDMs derived from wild-type or the *gp91phox^-/-^*mice were infected with *Francisella tularensis* LVS or the Δ*oxyR* mutant at a multiplicity of infection of 50 for 24 hours. (**A, B**) Levels of GBP5 and GBP2 in wild-type and (**C, D**) in *gp91phox^-/-^* BMDMs were assessed by SDS-PAGE followed by Western blot analysis. β-Actin was used as a loading control. Representative immunoblots from three independent experiments are shown. Band intensities (**B and D**) were determined by densitometric analysis, and data from three independent experiments were combined and analyzed using one-way ANOVA. *P* values are indicated (\*\*\**P* < 0.001).

### Enhanced inflammasome activation observed in BMDMs infected with the Δ*oxyR* mutant is ROS-dependent

We investigated whether the enhanced inflammasome activation observed in BMDMs infected with the Δ*oxyR* mutant is ROS-dependent. We infected BMDMs derived from the wild-type and *gp91phox^-/-^* mice with *F. tularensis* LVS and the Δ*oxyR* mutant. The macrophages were lysed and analyzed by SDS-PAGE and western blot to detect bioactive caspase-1 and IL-1β. Significantly higher levels of bioactive caspase-1 and IL-1β were detected in BMDMs derived from wild-type mice infected with the Δ*oxyR* mutant (Fig. 5A and B). However, no detectable bands of bioactive caspase-1 or IL-1β were observed in *gp91phox^-/-^* BMDMs infected either with *F. tularensis* LVS or the Δ*oxyR* mutant (Fig. 5C and D). These results indicate that the enhanced inflammasome activation in BMDMs infected with the Δ*oxyR* mutant is ROS-dependent.

**FIG 5:**
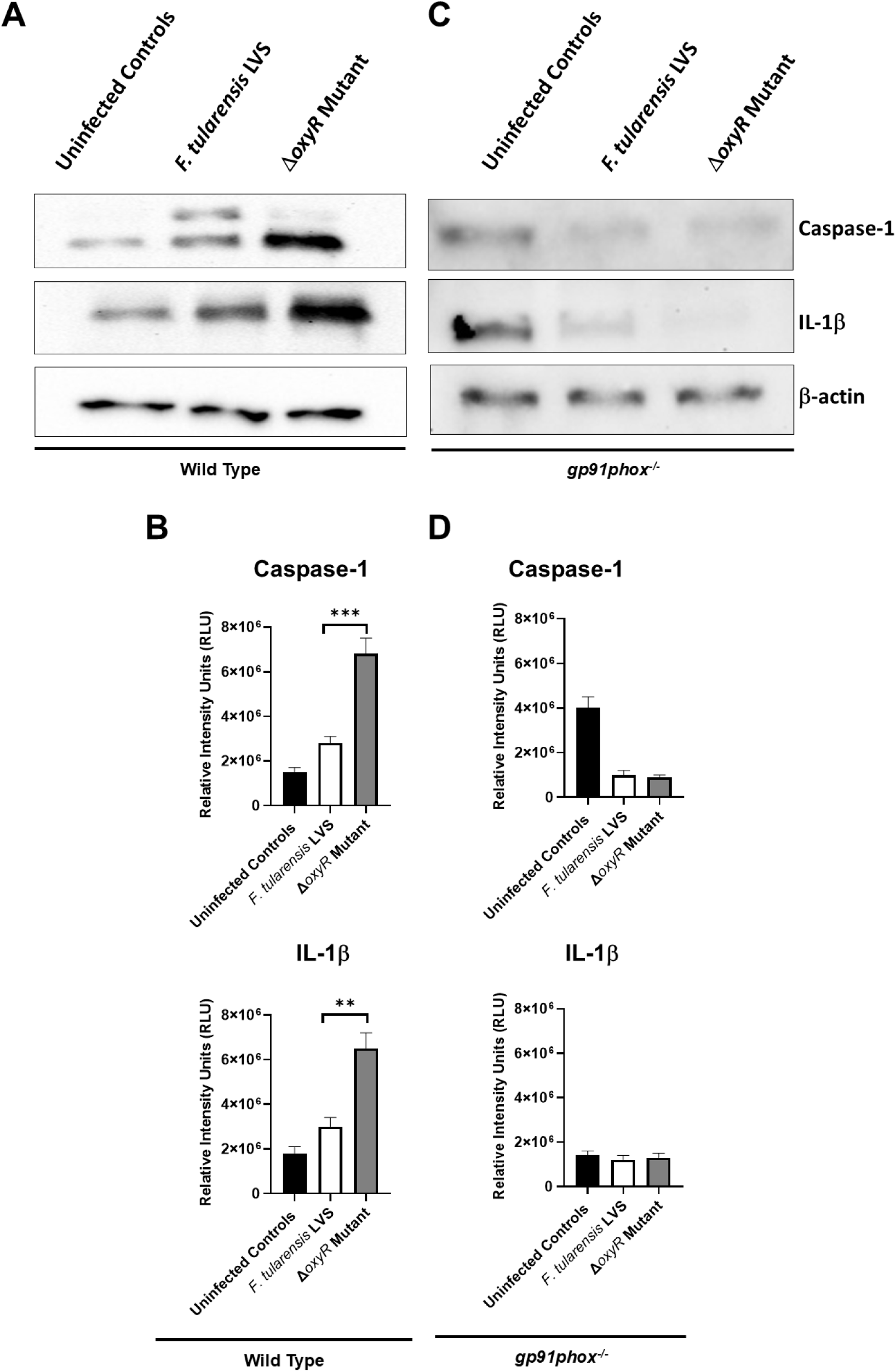
Enhanced inflammasome activation observed in BMDMs infected with the ΔoxyR mutant is ROS-dependent. BMDMs derived from wild-type **(A, B)** or the *gp91phox^-/-^* mice (**C, D**) were infected with *Francisella tularensis* LVS or the Δ*oxyR* mutant at a multiplicity of infection of 50 for 24 hours. (**A, C**) Levels of bioactive caspase-1 and IL-1β were assessed by SDS-PAGE followed by Western blot analysis. β-Actin was used as a loading control. Representative immunoblots from three independent experiments are shown. **(B, D)** Quantification of bioactive caspase-1 and IL-1β levels. Band intensities were determined by densitometric analysis, and data from three independent experiments were combined and analyzed using one-way ANOVA. *P* values are indicated (\*\**P* < 0.01, \*\*\**P* < 0.001).

### Inflammasome-mediated immune response can be altered by modulating the macrophage redox environment

Since our preceding results suggest that infection of BMDMs with the Δ*oxyR* mutant results in enhanced activation of bioactive forms of caspase-1 and IL-1β suggesting that the antioxidant enzymes of *F. tularensis* may have an essential role in suppression of Aim2 inflammasome-mediated immune response, we next investigated if similarly elevated levels of IL-1β are also observed in BMDMs infected with mutants of *F. tularensis* LVS deficient in primary antioxidant enzymes. BMDMs were infected with *F. tularensis* LVS or the *emrA1, sodB*Δ*sodC,* and Δ*katG* mutants. The Δ*oxyR* mutant was also included in this study. The macrophage cell lysates were analyzed for bioactive IL-1β 24 hours post-infection by western blot analysis. BMDMs infected with the *emrA1*, *sodB*Δs*odC,* and Δ*oxyR* mutants exhibited significantly enhanced levels of bioactive IL-1β when compared with the wild-type *F. tularensis* LVS. (Fig. 6A and B). Collectively, these results suggest that the loss of antioxidant enzymes leads to increased activation of the inflammasome, which in turn results in enhanced levels of bioactive IL-1β.

**FIG 6:**
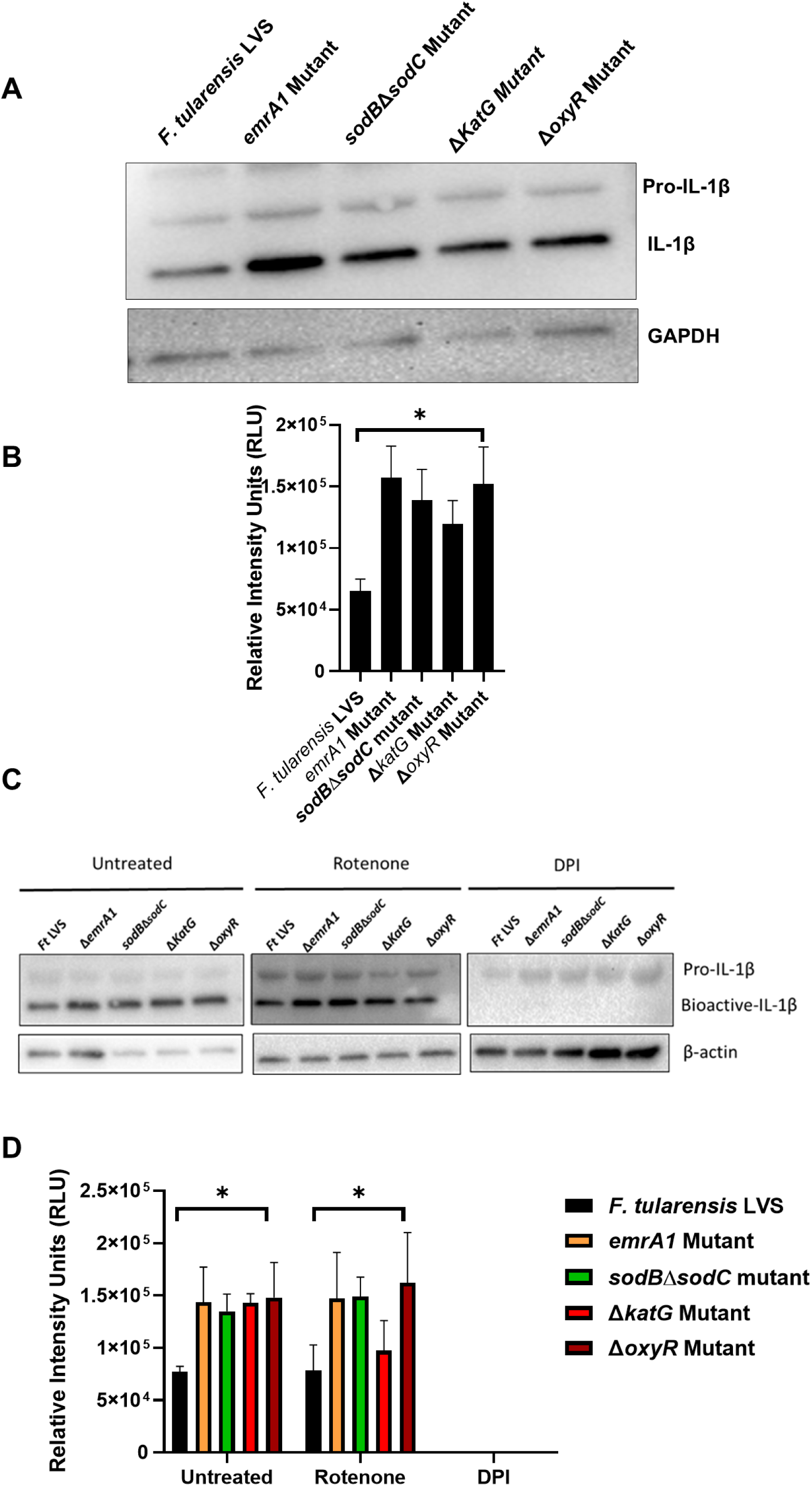
Inflammasome-mediated immune response can be altered by modulating the macrophage redox environment. BMDMs derived from wild type were infected with *Francisella tularensis* LVS or the indicated mutants for 24 hours. (**A**) Levels of bioactive IL-1β were assessed by SDS-PAGE followed by Western blot analysis. GAPDH was used as a loading control. Representative immunoblots from three independent experiments are shown. **(B)** Quantification of bioactive IL-1β levels. Band intensities were determined by densitometric analysis, and data from three independent experiments were combined and analyzed using one-way ANOVA. *P* values are indicated (\**P* < 0.05). **(C)** BMDMs derived from wild-type mice were treated with rotenone or diphenyleneiodonium chloride (DPI) as described in the Materials and Methods section and infected with *Francisella tularensis* LVS or the indicated mutants for 24 hours. Levels of bioactive IL-1β were assessed by SDS-PAGE followed by Western blot analysis. β-Actin was used as a loading control. Representative immunoblots from three independent experiments are shown. **(B)** Quantification of bioactive IL-1β levels. Band intensities were determined by densitometric analysis, and data from three independent experiments were combined and analyzed using one-way ANOVA. *P* values are indicated (\**P* < 0.05).

To further address the precise role of the antioxidant enzymes of *F. tularensis* LVS, we conducted experiments designed to trigger the production of ROS in infected BMDMs and investigated the impact of enhanced ROS production on bioactive IL-1β levels. The BMDMs were left untreated, treated with the complex I inhibitor rotenone, which induces mitochondrial ROS production, or treated with DPI, an inhibitor of NADPH-oxidase-dependent ROS. As observed earlier, enhanced levels of bioactive IL-1β were detected in BMDMs infected with mutants of *F. tularensis* deficient in antioxidant enzymes (Fig. 6C). However, when BMDMs were treated with rotenone, the bioactive levels of IL-1β did not change in BMDMs infected with *F.tularensis* LVS and remained similar to those observed for the untreated BMDMs. On the other hand, the levels of bioactive IL-1β remained elevated in BMDMs infected with the antioxidant enzyme-deficient mutants, similar to their untreated counterparts. It was further observed that when the BMDMs were treated with ROS inhibitor DPI, the bioactive IL-1β was neither observed in BMDMs infected with *F. tularensis* LVS nor in those infected with mutants deficient in antioxidant enzymes (Fig. 6C and D). These results were similar to those observed for the *gp91phox^-/-^* BMDMs infected with the Δ*oxyR* mutant (Fig. 5C and D). Collectively, these results indicate that ROS play a major role in the induction of Aim2-dependent IL-1β production, and the antioxidant enzymes play an essential role in suppressing this response primarily by scavenging the ROS.

## DISCUSSION

The primary virulence mechanism of *F. tularensis* identified to date involves suppression of innate immune responses, including inflammasome activation (17, 18). This immunosuppression permits uncontrolled bacterial replication, extensive tissue damage, and progression to a sepsis-like syndrome and high mortality. Activation of the Aim2 inflammasome during *F. novicida* infection requires two distinct signaling pathways. Upon phagosomal escape, trace amounts of bacterial DNA are detected by the cGAS–STING pathway, inducing type I IFN and IRFs that promote the expression of GBPs (26). The GBPs bind to bacterial surfaces and induce lysis, releasing sufficient DNA to prime and activate the Aim2 inflammasome (14). In parallel, macrophage-derived microbicidal oxidants also cause bacterial lysis, further increasing cytosolic DNA (15). Aim2 inflammasome assembly leads to caspase-1 activation, cleavage of Gsdmd, and maturation of IL-1β and IL-18, which are released through Gsdmd-formed membrane pores.

The role and regulation of the Aim2 inflammasome during tularemia caused by *F. tularensis* remain incompletely understood. Our previous work showed that *F. tularensis* SchuS4 and *F. tularensis* LVS induce minimal IL-1β and IL-18 production and either suppress or weakly activate the Aim2 inflammasome in infected macrophages (17, 18). We hypothesized that these differences arise from distinct redox environments generated during infection, attributable to the superior oxidant-scavenging capacity of *F. tularensis*. To test this, we used a Δ*oxyR* mutant of *F. tularensis* LVS, generated in our laboratory. OxyR is the master regulator of oxidative stress responses in *F. tularensis*, controlling key antioxidant enzymes SodB, KatG, and AhpC, as well as more than 100 additional oxidative stress-related genes (8). Because *F. tularensis* lacks a SoxR homolog, OxyR is its sole oxidative stress regulator. The ΔoxyR mutant is highly sensitive to oxidative stress and fails to control the intracellular redox environment of infected macrophages (8).

Our results show that *F. tularensis* LVS induces weak or suppressed Aim2 inflammasome activation in BMDMs, as evidenced by low levels of active caspase-1 and IL-1β. In contrast, infection with the Δ*oxyR* mutant significantly increased Aim2-dependent caspase-1 activation and IL-1β production. Although cGAS-STING signaling is essential for Aim2 activation during *F. novicida* infection (14, 26), STING deficiency did not affect inflammasome activation in BMDMs infected with the Δ*oxyR* mutant. Similarly, IRF1 induction occurred independently of STING, indicating a distinct upstream mechanism.

Our results demonstrate that increased oxidative stress during Δ*oxyR* mutant infection activates STAT1, which upregulates IRF1 and GBPs 2 and 5, promoting bacterial lysis and Aim2 activation. It was observed that STAT1 phosphorylation and downstream IRF1 and GBP expression were ROS-dependent and markedly reduced in *gp91phox*^-/-^ macrophages. Correspondingly, Aim2 activation, caspase-1 cleavage, and IL-1β release were diminished in these cells, highlighting ROS as a central driver of inflammasome activation in this context. This notion is further cemented by the evidence that enhanced levels of bioactive IL-1β were also observed in BMDMs infected with mutants deficient in *emrA1*, which is highly sensitive to oxidative stress due to its inability to secrete KatG and SodB (20)*, sodB*Δ*sodC,* and Δ*katG* mutants. Further, the treatment of BMDMs with rotenone, which enhances the accumulation of mitochondrial ROS (27) results in sustained elevated bioactive levels of bioactive IL-1β in BMDMs infected with the antioxidant enzyme-deficient mutants of *F. tularensis* LVS than those observed in untreated BMDMs. Due to fully functional antioxidant enzymes in the wild type *F. tularensis* LVS, rotenone treatment does not have any effect in terms of an increase in bioactive IL-1β levels. However, this effect is more pronounced in BMDMs infected with the mutants deficient in antioxidant enzymes. Furthermore, complete abolition of ROS by treating macrophages with ROS inhibitor DPI (28) abrogates bioactive IL-1β in the wild type as well as mutant infected BMDMs, a feature similar to that observed for the *gp91phox^-/-^* BMDMs. These results indicate that the redox environment of macrophages following infection plays an important role in the activation of Aim2 inflammasome-dependent cytokines in response to *F. tularensis* LVS. Further, the results demonstrate that *F. tularensis* exploits its robust antioxidant defenses to neutralize macrophage-derived ROS, thereby suppressing redox-dependent Aim2 inflammasome activation and proinflammatory cytokine production. Collectively, this study uncovers a critical virulence mechanism and the complex immune evasion strategy of *F. tularensis*. Our findings reveal a novel role for bacterial antioxidant systems in dampening host innate immunity.

## ACKNOWLEDGEMENTS

This work was supported by National Institutes of Health Grants R15AI107698 (MM). The funders had no role in study design, data collection and analysis, decision to publish, or manuscript preparation. No financial conflicts of interest exist regarding the contents of the manuscript and its authors.

## REFERENCES

1. Francis E. 1925. Tularemia. J Am Med Assoc 84:1243–1250.

2. Dennis DT, Inglesby T v., Henderson DA, Bartlett JG, Ascher MS, Eitzen E, Fine AD, Friedlander AM, Hauer J, Layton M, Lillibridge SR, McDade JE, Osterholm MT, ÖToole T, Parker G, Perl TM, Russell PK, Tonat K. 2001. Tularemia as a biological weapon: Medical and public health management. J Am Med Assoc 285:2763–2773.

3. Telford SR, Goethert HK. 2020. Ecology of *Francisella tularensis*. Annu Rev Entomol 65:351–372.

4. Kingry LC, Petersen JM. 2014. Comparative review of Francisella tularensis and Francisella novicida. Front Cell Infect Microbiol 4.

5. Melillo AA, Mahawar M, Sellati TJ, Malik M, Metzger DW, Melendez JA, Bakshi CS. 2009. Identification of Francisella tularensis Live Vaccine Strain CuZn Superoxide Dismutase as Critical for Resistance to Extracellularly Generated Reactive Oxygen Species. J Bacteriol 191:6447–6456.

6. Bakshi CS, Malik M, Regan K, Melendez JA, Metzger DW, Pavlov VM, Sellati TJ. 2006. Superoxide Dismutase B Gene (sodB)-Deficient Mutants of Francisella tularensis Demonstrate Hypersensitivity to Oxidative Stress and Attenuated Virulence. J Bacteriol 188:6443–6448.

7. Alharbi A, Rabadi SM, Alqahtani M, Marghani D, Worden M, Ma Z, Malik M, Bakshi CS. 2019. Role of peroxiredoxin of the AhpC/TSA family in antioxidant defense mechanisms of Francisella tularensis. PLoS One 14:e0213699.

8. Ma Z, Russo VC, Rabadi SM, Jen Y, Catlett S V., Bakshi CS, Malik M. 2016. Elucidation of a mechanism of oxidative stress regulation in Francisella tularensis live vaccine strain. Mol Microbiol 101:856–878.

9. Lindgren H, Shen H, Zingmark C, Golovliov I, Conlan W, Sjöstedt A. 2007. Resistance of Francisella tularensis strains against reactive nitrogen and oxygen species with special reference to the role of KatG. Infect Immun 75:1303–1309.

10. Zhang J-R, Pechous RD, Honn M, Lindgren H, Bharath GK, Sjöstedt A. 2017. Lack of OxyR and KatG Results in Extreme Susceptibility of Francisella tularensis LVS to Oxidative Stress and Marked Attenuation In vivo. Frontiers in Cellular and Infection Microbiology | www.frontiersin.org 1:14.

11. Zhu Q, Man SM, Karki R, Malireddi RKS, Kanneganti T-D. 2018. Detrimental Type I Interferon Signaling Dominates Protective AIM2 Inflammasome Responses during Francisella novicida Infection. Cell Rep 22:3168–3174.

12. Henry T, Monack DM. 2007. Activation of the inflammasome upon Francisella tularensis infection: Interplay of innate immune pathways and virulence factors. Cell Microbiol 10.1111/j.1462-5822.2007.01022.x.

13. Henry T, Brotcke A, Weiss DS, Thompson LJ, Monack DM. 2007. Type I interferon signaling is required for activation of the inflammasome during Francisella infection. J Exp Med 204:987–994.

14. Meunier E, Wallet P, Dreier RF, Costanzo S, Anton L, Rühl S, Dussurgey S, Dick MS, Kistner A, Rigard M, Degrandi D, Pfeffer K, Yamamoto M, Henry T, Broz P. 2015. Guanylate-binding proteins promote activation of the AIM2 inflammasome during infection with Francisella novicida. Nat Immunol 16:476–84.

15. Banerjee I, Behl B, Mendonca M, Shrivastava G, Russo AJ, Menoret A, Ghosh A, Vella AT, Vanaja SK, Sarkar SN, Fitzgerald KA, Rathinam VAK. 2018. Gasdermin D Restrains Type I Interferon Response to Cytosolic DNA by Disrupting Ionic Homeostasis. Immunity 49:413–426.e5.

16. Crane DD, Bauler TJ, Wehrly TD, Bosio CM. 2014. Mitochondrial ROS potentiates indirect activation of the AIM2 inflammasome. Front Microbiol 5.

17. Dotson RJ, Rabadi SM, Westcott EL, Bradley S, Catlett S V., Banik S, Harton JA, Bakshi CS, Malik M. 2013. Repression of Inflammasome by Francisella tularensis during Early Stages of Infection. Journal of Biological Chemistry 288:23844–23857.

18. Alqahtani M, Ma Z, Miller J, Yu J, Malik M, Bakshi CS. 2023. Comparative analysis of absent in melanoma 2-inflammasome activation in Francisella tularensis and Francisella novicida. Front Microbiol 14.

19. Bakshi CS, Mahawar M, Melendez JA, Sellati TJ, Malik M, Metzger DW, Melillo AA, Mahawar M, Sellati TJ, Malik M, Metzger DW, Melendez JA, Bakshi CS. 2009. Identification of Francisella tularensis Live Vaccine Strain CuZn Superoxide Dismutase as Critical for Resistance to Extracellularly Generated Reactive Oxygen Species. J Bacteriol 10.1128/jb.00534-09.

20. Ma Z, Banik S, Rane H, Mora VT, Rabadi SM, Doyle CR, Thanassi DG, Bakshi CS, Malik M. 2014. EmrA1 membrane fusion protein of Francisella tularensisLVS is required for resistance to oxidative stress, intramacrophage survival and virulence in mice. Mol Microbiol 91:976–995.

21. Storek KM, Gertsvolf NA, Ohlson MB, Monack DM. 2015. cGAS and Ifi204 cooperate to produce type I IFNs in response to Francisella infection. J Immunol 194:3236–3245.

22. Almine JF, O’Hare CAJ, Dunphy G, Haga IR, Naik RJ, Atrih A, Connolly DJ, Taylor J, Kelsall IR, Bowie AG, Beard PM, Unterholzner L. 2017. IFI16 and cGAS cooperate in the activation of STING during DNA sensing in human keratinocytes. Nat Commun 8:1–15.

23. Henry T, Brotcke A, Weiss DS, Thompson LJ, Monack DM. 2007. Type I interferon signaling is required for activation of the inflammasome during Francisella infection. J Exp Med 204:987–994.

24. Ramsauer K, Farlik M, Zupkovitz G, Seiser C, Kröger A, Hauser H, Decker T. 2007. Distinct modes of action applied by transcription factors STAT1 and IRF1 to initiate transcription of the IFN-γ-inducible *gbp2* gene. Proceedings of the National Academy of Sciences 104:2849–2854.

25. Soond SM, Townsend PA, Barry SP, Knight RA, Latchman DS, Stephanou A. 2008. ERK and the F-box Protein βTRCP Target STAT1 for Degradation. Journal of Biological Chemistry 283:16077–16083.

26. Man SM, Karki R, Malireddi RKS, Neale G, Vogel P, Yamamoto M, Lamkanfi M, Kanneganti T. 2015. The transcription factor IRF1 and guanylate-binding proteins target activation of the AIM2 inflammasome by Francisella infection. Nat Immunol 16:1–11.

27. Li N, Ragheb K, Lawler G, Sturgis J, Rajwa B, Melendez JA, Robinson JP. 2003. Mitochondrial Complex I Inhibitor Rotenone Induces Apoptosis through Enhancing Mitochondrial Reactive Oxygen Species Production. Journal of Biological Chemistry 278:8516–8525.

28. Buck A, Sanchez Klose FP, Venkatakrishnan V, Khamzeh A, Dahlgren C, Christenson K, Bylund J. 2019. DPI Selectively Inhibits Intracellular NADPH Oxidase Activity in Human Neutrophils. Immunohorizons 3:488–497.

